# Evolving Notch polyQ tracts reveal possible solenoid interference elements

**DOI:** 10.1101/079038

**Authors:** Albert J. Erives

## Abstract

Polyglutamine (polyQ) tracts in regulatory proteins are extremely polymorphic. As functional elements under selection for length, triplet repeats are prone to DNA replication slippage and indel mutations. Many polyQ tracts are also embedded within intrinsically disordered domains, which are less constrained, fast evolving, and difficult to characterize. To identify structural principles underlying polyQ tracts in disordered regulatory domains, here I analyze deep evolution of metazoan Notch polyQ tracts, which can generate alleles causing developmental and neurogenic defects. I show that Notch features polyQ tract turnover that is restricted to a discrete number of conserved “polyQ insertion slots”. Notch polyQ insertion slots are: (*i*) identifiable by an amphipathic “slot leader” motif; (*ii*) conserved as an intact C-terminal array in a 1-to-1 relationship with the N-terminal solenoid-forming ankyrin repeats (ARs); and (*iii*) enriched in carboxamide residues (Q/N), whose sidechains feature dual hydrogen bond donor and acceptor atoms. Correspondingly, the terminal loop and β-strand of each AR feature conserved carboxamide residues, which would be susceptible to folding interference by hydrogen bonding with residues outside the ARs. I thus suggest that Notch polyQ insertion slots constitute an array of AR interference elements (ARIEs). Notch ARIEs would dynamically compete with the delicate serial folding induced by adjacent ARs. Huntingtin, which harbors solenoid-forming HEAT repeats, also possesses a similar number of polyQ insertion slots. These results strongly suggest that intrinsically disordered interference arrays featuring carboxamide and polyQ enrichment are coupled proteodynamic modulators of solenoids.

**SIGNIFICANCE:** Neurodegenerative disorders are often caused by expanded polyglutamine (polyQ) tracts embedded in the disordered regions of regulatory proteins, which are difficult to characterize structurally. To identify functional principles underlying polyQ tracts in disordered regulatory domains, I analyze evolution of the Notch protein, which can generate polyQ-related alleles causing neurodevelopmental defects. I show that Notch evolves polyQ tracts that come and go in a few conserved “polyQ insertion slots”. Several features suggest these slots are ankyrin repeat (AR) interference elements, which dynamically compete with the delicate solenoid formed by Notch. Huntingtin, whose polyQ expansions causes Huntington’s Disease in humans, also has solenoid-forming modules and polyQ insertion slots, suggesting a common architectural principle underlies solenoid-forming polyQ-rich proteins.

## INTRODUCTION

Polyglutamine (polyQ) tracts are functional features of many conserved transcriptional regulators and allow for complex conformational dynamics and interactions with other polyQ factors (1–9). A prominent polyQ tract was first identified in the neurogenic gene *Notch* (10). The Notch protein is central to a signaling pathway guiding patterning and cell fate decisions during metazoan development (11). This polyQ tract is embedded in the Notch intracellular domain (NICD), which when cleaved leads to nuclear import and activation via DNA-bound CSL proteins. This polyQ tract is highly polymorphic in *Drosophila melanogaster* and can generate new alleles that cause developmental and neurogenic defects (7). All of the identified polymorphic alleles are specific to *Drosophila melanogaster* because this same tract is uniquely configured in most other *Drosophila* species as determined by the placement of an intervening histidine residue and the underlying CAX triplet nucleotide repeat pattern (7).

The single polyQ tract of *Drosophila* Notch proteins is embedded in a much larger intrinsically disordered protein (IDP) region, which is common to many transcriptional activators and co-activators, including NICD, TBP, and many of the Mediator co-activator subunits (12, 13). It is also common to large scaffolding proteins that serve as signal integration platforms that are regulated by multiple protein-protein interactions. On such example is Huntingtin (Htt), which is conserved in humans, flies (14), and non-metazoan eukaryotes such as social slime molds (15). Polyglutamine (polyQ) expansions in Htt (1, 8, 9, 16, 17), TBP (18, 19), and other conserved loci (1) in humans underlie various progressive neurodegenerative diseases.

IDP regions of regulator and scaffolding proteins provide conformational flexibility and allow the formation of transient regulator complexes under specific conditions (12, 13). IDPs form random coils or molten globules with very little protein secondary structure and this makes them exceedingly difficult to study biophysically. As such, these regions are typically removed in proteins subjected to structural studies. These regions are also difficult to align with orthologous sequences encoded in other genomes, and are a major impediment to accurate gene annotation in whole genome sequence assemblies. Furthermore, short read assemblies are intractable at loci encoding lengthy polyQ tracts.

Although some eukaryotes, such as the amoebozoan slime mold *Dictyostelium*, exhibit genome-wide trends in polyQ content (20–22), in general encoded polyQ tracts of conserved regulators expand and contract in a locus-specific manner. The N-terminal polyQ tract in human Htt is highly polymorphic and is expanded relative to early branching primates (*e.g*., tarsiers), and other mammals (*e.g*., mouse) (Fig. 1a, **top**). However, the *Drosophila* Htt ortholog features only a single glutamine residue at this position (Fig. 1a, **top**). Nonetheless, a different region of *Drosophila* Htt features a prominent polyQ tract, which is absent in vertebrates, including humans, despite the region of its appearance being conserved (Fig. 1a, **bottom**). Both of these polyQ “insertion slots” are preceded by a short leader motif that is predicted to form an amphipathic helix. In general, polyQ tracts evolve in an evolutionary turnover process, the study of which can lead to biophysical insights not immediately accessible via biochemical characterization.

**Figure 1.**
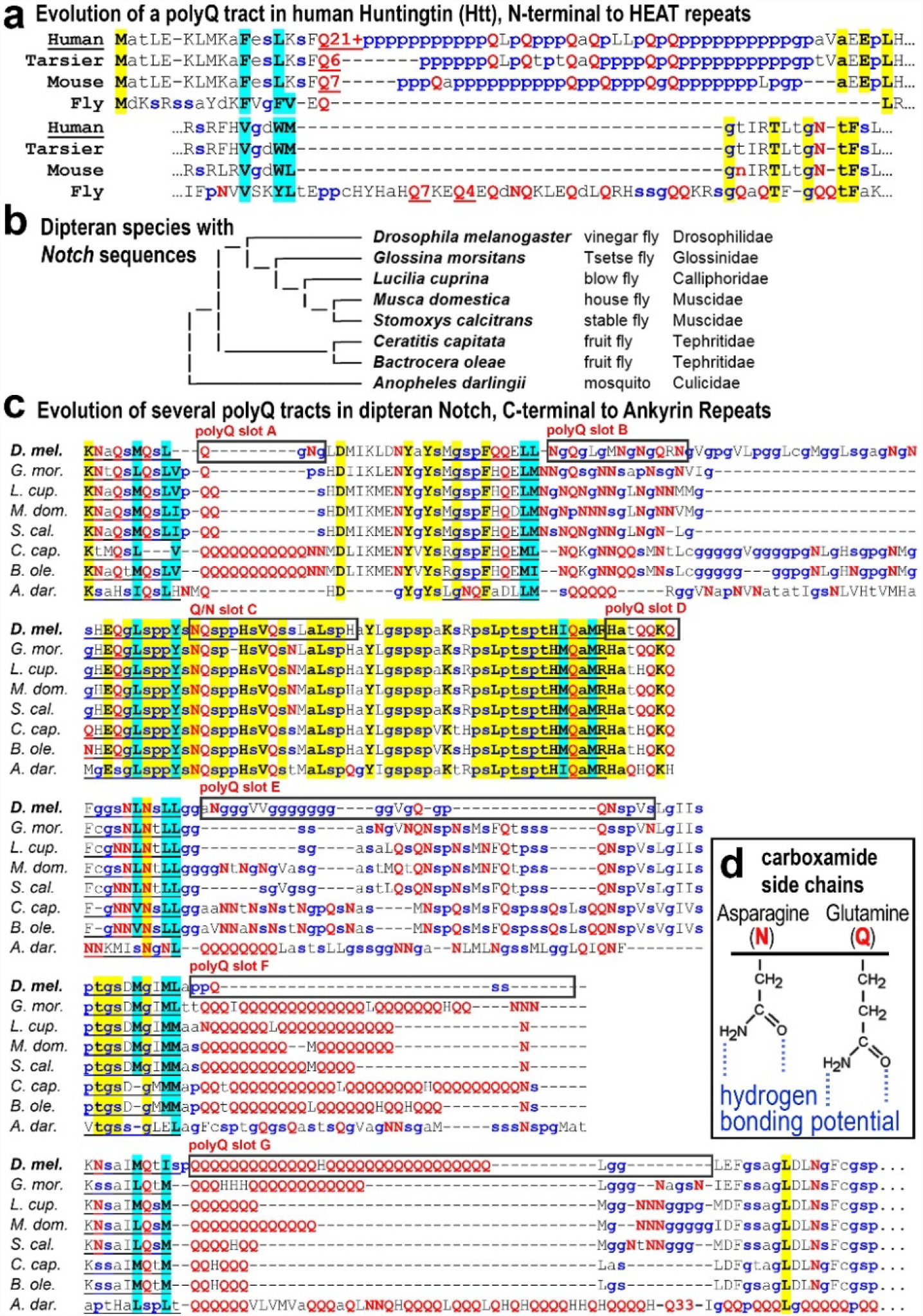
PolyQ tracts evolve in well-defined slots in Htt and Notch proteins. **(a)** Shown is the N-terminal region of Huntingtin (Htt) from humans, tarsier, mouse, and *Drosophila*. This region evolved a polyglutamine (polyQ) tract at a well-defined location in mammals, which later expanded in the evolution of humans, and is still evolving. While fly Htt is missing the N-terminal polyQ tract, it features a separate tract elsewhere in the protein as shown. Small amino acid residues that are secondary structure breakers are highlighted in blue. Glutamine residues are highlighted in red. Hydrophobic amino acid residues preceding the polyQ tract insertion site are highlighted in cyan. Conserved residues are highlighted in yellow. **(b)** Shown is an evolutionary tree of various dipteran genera for which Notch sequences were analyzed in this study. The six different families to which these species belong are listed. **(c)** Shown is the polyQ tract regions of fly Notch proteins. Several sites (lettered slots) have independently evolved a polyQ tract in different fly genera. Most slots are preceded by a leader motif sequence, even when a polyQ tract is not present. **(d)** Inset shows the high hydrogen bonding potential of carboxamide amino acid side chains.

To better understand the biochemical constraints governing polyQ evolution, I considered the long polyQ tract in the Notch intracellular domain in *Drosophila*. We previously found that this polyQ tract is highly unstable and variable within the *Drosophila melanogaster* population (7). This tract has been continuously evolving since the *Drosophila* radiation with different species exhibiting distinct polyQ tract configurations within the large IDP region of NICD (7). Here I analyzed Notch proteins from diverse dipterans (flies) and discovered a previously unknown level of Notch polyQ evolution that is functionally revealing about the Notch intrinsically ordered region and its relation to the adjacent solenoid-forming ankyrin repeats (AR). I show that a polyQ slot insertion model provides a powerful interpretation of the functional interactions between polyQ tracts constrained to a limited number of slots in solenoid-type proteins such as the Notch intracellular domain and Huntingtin. The polyQ insertion model can be informed by use of the fast-evolving dipteran orthologs of human disease-related proteins featuring expanded polyQ tracts.

## RESULTS

I define “polyQ insertion slots” as specific positions in a protein where polyQ tracts are favored to occur during evolution. Strong support for a polyQ insertion model of protein evolution could include (*i*) evidence of a non-random or constrained distribution of polyQ tracts across a set of diverged orthologs, (*ii*) evidence of a polyQ turnover process whereby polyQ tracts have been lost at one position and gained at another, (*iii*) identification of structural or peptide sequence motifs serving as polyQ slot leaders or trailers, and (*iv*) evidence of a functional relationship of polyQ slots to other functional domains within the protein or within a protein interactor.

To identify polyQ insertion slots in the Notch intracellular domain, I analyzed predicted Notch proteins from eight different genera in six different families of Diptera (Fig. 1b). I find that polyQ tracts in dipteran Notch proteins have evolved in seven different well-defined polyQ insertion slots (Fig. 1c). Each slots is preceded by a conserved polyQ slot leader motif similar to the Htt amphipathic helix, further confirming the utility of the polyQ insertion slot model. Even in the absence of polyQ tracts, slot leaders precede a region enriched in the carboxamide residues, asparagine (N) and glutamine (Q), whose sidechains feature hydrogen bond donor and acceptor groups (Fig. 1d). These same slot regions are also enriched in the small amino acid residues that are secondary structure breakers (“g”, “p”, and “s”).

Using the polyQ slot leader organizational model of the IDP region of dipteran NICD and a few reliably homologous slot leader motifs, I identified seven slots in the human Notch1 protein, as shown by an alignment to *Drosophila* NICD (Fig. 2a). As in dipteran Notch proteins, these slots in human Notch1 are also enriched in both carboxamide residues and small secondary structure breakers that lead to IDPs. Thus, carboxamide-enriched IDPs with slot leaders likely represent a deeper organizational principle underlying many polyQ-rich regulators. I demonstrate the utility of this point as follows.

**Figure 2.**
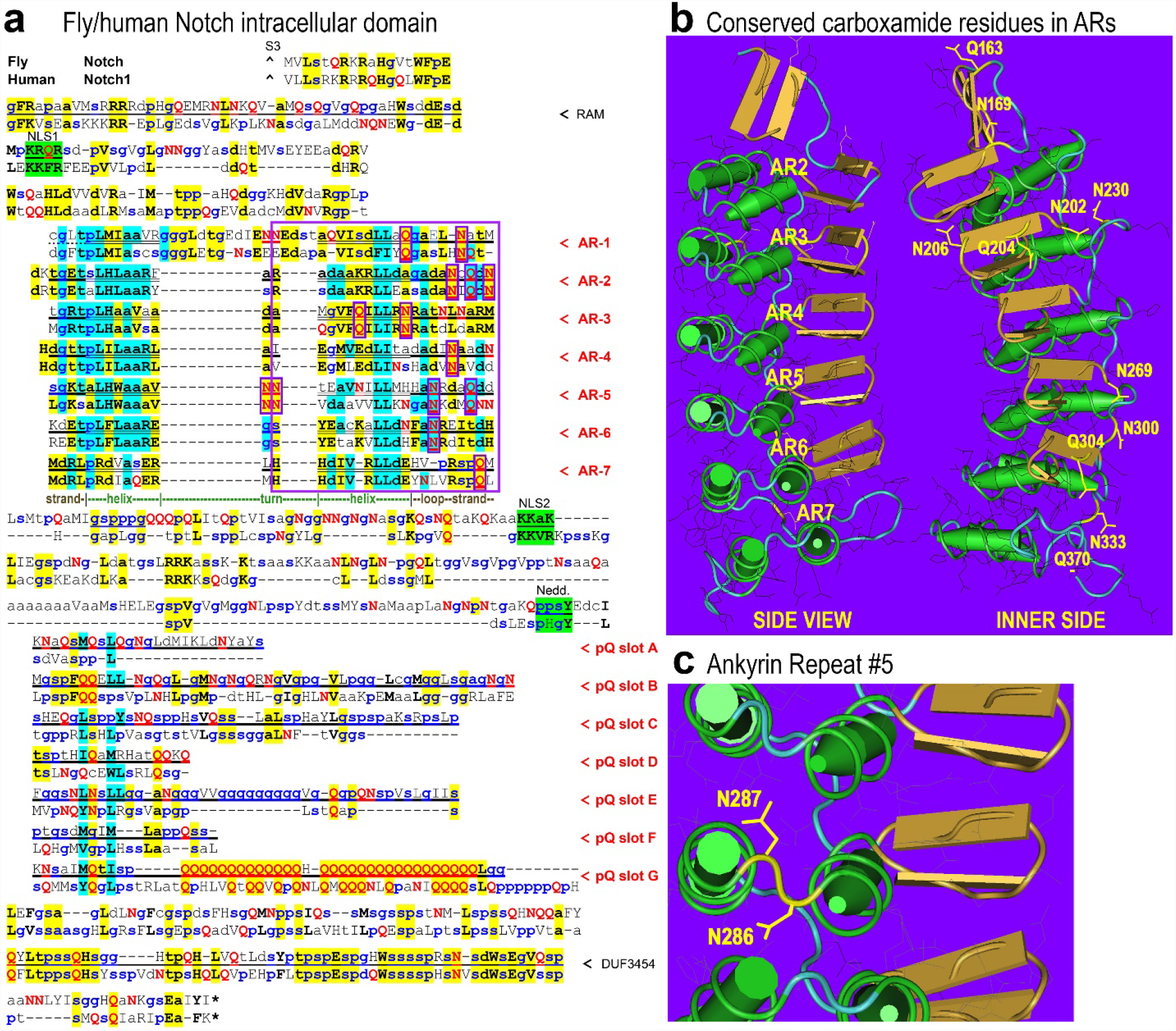
Notch polyQ slots are conserved IDP interference elements of ankyrin repeats. **(a)** Shown is an alignment of *Drosophila* Notch and human Notch1 intracellular domains (NICD) beginning at the S3 cleavage site (caret). The CSL/RBPJ-associated molecule (RAM) region is located at the N-terminus of NICD. Two nuclear localization signals (green with wavy underlining) flank the seven ankyrin repeats (ARs). The 33 residues of each AR features a helix-turn-helix peptide motif (helical regions in double underlining). This is followed by a conserved neddylation site and the six or seven polyQ insertion slots each starting with a polyQ slot leader sequence motif. A conserved domain of unknown function (DUF3454) follows the polyQ insertion slots. The polyQ slots fill up the region that is intrinsically disordered, but some slots are more conserved in sequence than others, such as slot-B, slot-D, and slot-G. **(b)** The crystal structure of the human AR2–AR7 region is annotated to show the carboxamide residues that are conserved between fly Notch and human Notch1. These conserved Q/N residues and others not conserved are featured in the terminal loop and strand elements of each ankyrin repeat. **(c)** Shown is a close-up of ankyrin repeat 5 and the conserved double asparagines in the turn element.

The seven N-terminal ankyrin repeats (ARs) in NICD form a solenoid-like domain, which fold by delicate intra-repeat interactions. Given the approximate 1-to-1 conservation of ARs to the IDP repeats with polyQ slots, it is possible that the IDP array is a kinetically active ensemble on one side of the linear AR solenoid. To investigate whether the polyQ IDP slots could be proteodynamic modulators of ARs, I analyzed the AR structural elements for carboxamide residues that could be hydrogen bonding with the carboxamide and polyQ rich IDPs. I find that the terminal halves of ARs are >10-fold enriched (13-to-1) in conserved carboxamide residues relative to the first half (**see boxed residues in** Fig. 2a, **and labeled residues in** Fig. 2b). Additional non-conserved carboxamide residues are also enriched in the terminal halves of ARs of both species, particularly in the terminal loop and strand elements (**red Q’s and N’s in** Fig. 2a).

The terminal β-strand of each AR forms a hairpin with the starting β-strand of the next AR, and is important for induced propagation of solenoid formation (23). I thus propose that the conserved array of seven polyQ insertion slots, which feature polyQ and/or carboxamide enriched IDPs, constitute AR interference elements (ARIEs). A pair of adjacent asparagines are also conserved in AR-5, which could be available for hydrogen bonding interactions in unfolded Notch AR solenoids (Fig. 2c). Furthermore, the approximate 1-to-1 conservation of ARs to ARIEs in Notch suggest that ARIEs are proteodynamic modulators can be associated with specific solenoid repeats or pairs of repeats. The proteodynamic ARIEs array would attenuate AR solenoid formation via kinetic coupling and constitute a critical aspect of NICD regulation at multiple steps in the Notch signaling pathway.

Analogous to Notch and other AR-based solenoids, the proteins <u>H</u>tt, <u>E</u>F3, the regulatory <u>A</u> subunit of PP2A, and <u>T</u>OR1 (target of rapamycin) are solenoids based on HEAT repeats with Htt having three HEAT repeats (24). Thus, I further suggest that solenoid interference arrays featuring polyQ turnover dynamics are functionally-coupled components of solenoids. Solenoid folding is a delicate regulatory-prone process and is distinct from the folding seen in stable globular structures driven by hydrophobic packing and long-distance interactions. Thus, a folding funnel landscape view of NICD and Htt might be described by bi-stable minima involving a lower moat ARIEs-ARs interference state and a higher dimple solenoid state (25).

The conserved carboxamide residues of NICD occupy the inner side of the AR solenoid (Fig. 2b, **numbered residues on inner side**). This side of the AR solenoid corresponds to the interaction surface in transcriptional complexes with Mastermind and CSL (23, 26). Possibly, the polyQ richness of Mastermind (Mam/Maml) (4, 5, 27), a dedicated Notch-coactivator, functions as an AR solenoid chaperone by competing with the same solenoid surface entangled with the ARIEs array. Similarly, the Notch interacting protein Deltex features polyQ tracts in *Drosophila*, which has a longer polyQ tract than human Notch proteins, but not in human Deltex homologs (not shown). As such these functional intermolecular interactions would stem directly from AR-ARIE intramolecular coupling. Intermolecular ARIE-ARIE interactions in transcriptional NICD multimers (28, 29) may also relieve intramolecular inhibition of AR solenoid formation by the ARIEs array. Thus, polyQ expansions and contractions are likely to be under complex selection for their regulatory interactions with the adjacent solenoid repeats and their regulatory modulators.

Analysis of the evolutionary turnover of polyQ tracts in dipteran NICD moieties provides a novel perspective of the IDP region of NICD, and possibly completes the functional domain inventory of a key developmental signaling molecule (Fig. 3a). ARIEs are evident by evolutionary polyQ slot turnover and by the subsequently identified common leader motif resembling the amphipathic leader in Huntingtin. To see if additional peculiarities pertain to the NICD slot leaders, I derived an NICD-specific polyQ slot leader motif by taking the nine residue peptide leader sequences from *Drosophila* and *Stomoxys* slot leaders except for those from the hypothesized polyQ slot-C, for which polyQ insertions have not yet been seen (Fig. 3b). Not only is this a short amphipathic helix with hydrophobic residues on one side but there is typically at least one glutamine residue adjacent to the hydrophobic side (Fig. 3b). Thus, the leader motif itself may function to both display and interact with polyQ epitopes or carboxamide-rich IDPs in slots, or with the carboxamide rich terminal elements of associated ankyrin repeats.

**Figure 3.**
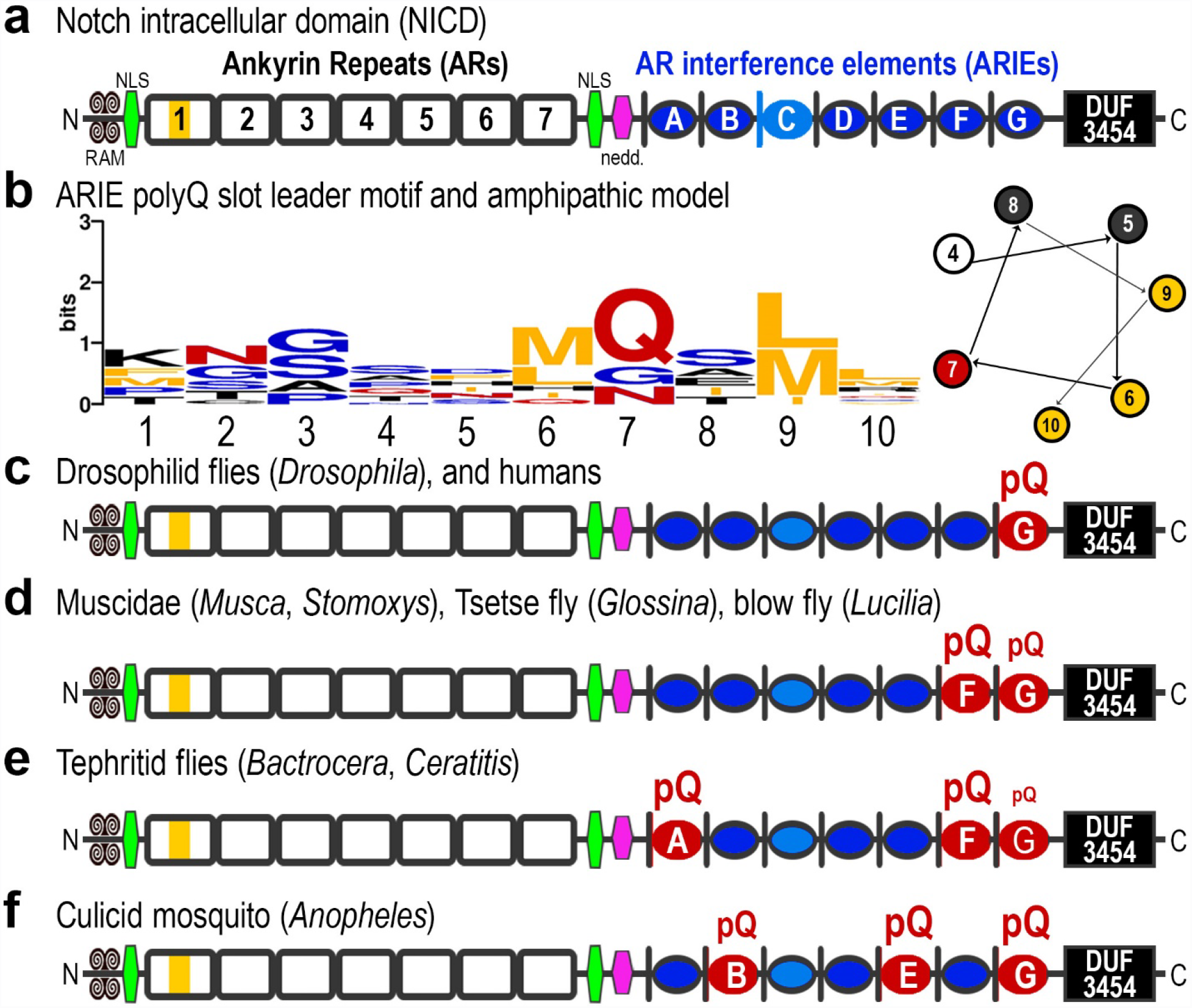
Notch AR interference elements (ARIEs) are present as a distinct array within NICD. **(a)** The polyQ slot leader motif defines 7 possible insertion sites, for which six have independently evolved polyQ tracts in distinct lineages of flies. The number of slots is also suggestive of an interference with the seven ankyrin repeats or the six pairs of adjacent repeats. **(b)** The leader sequences from *Drosophila* and *Stomoxys* polyQ insertion slots (except for slot C) were used to produce the motif logo shown. Also shown is an amphipathic helix starting with residue number four and showing how the hydrophobic amino acids at positions 6, 9, and 10 of the slot leader motif are segregated to one side of the helix (yellow circles). Interestingly, position 7 frequently features a single glutamine residue, which could interact with an adjacent polyQ tract. **(c)** *Drosophila* Notch and vertebrate Notch1 feature a polyQ tract in slot-G, although it is much more expanded in *Drosophila* than humans. **(d, e)** Unlike *Drosophila* Notch, the Notch proteins from several other fly genera, including *Musca*, *Stomoxys*, *Glossina*, *Lucilia*, *Bactrocera*, and *Ceratitis*, feature a more prominent polyQ tract in slot-F. These flies also have a much smaller polyQ tract in slot-G. The tephritid genera also feature a tract in slot-A, demonstrating that N-terminal slots can also accept polyQ tracts. **(f)** The Notch protein from *Anopheles darlingii* features polyQ tracts in slot-B and slot-E in addition to a very long tract in slot-G.

## DISCUSSION

Altogether, the dipteran Notch proteins reveal the evolutionary appearance of lengthy polyQ tracts in five of the seven ARIEs and constitute an evolutionary Rosetta Stone for understanding NICD and its disordered carboxamide interference elements (Fig. 3c–f). The single polyQ tract of *Drosophila melanogaster* is polymorphic and homologous to the single polyQ tract that is configured differently in other *Drosophila* species. However, order-wide patterns across multiple dipteran genera are needed to reveal the entire expanse within which polyQ turnover happens. Similar deep evolutionary profiling of polyQ turnover in other conserved regulators may reveal additional solenoid-modulating interference arrays. Thus fast evolving dipteran genera may provide an ideal evolutionary system for understanding mutant human proteins associated with proteotoxicity diseases.

These results also raise the likelihood of blind spots in our understanding of solenoids formed by ankyrin repeats in Notch and other AR solenoids, HEAT repeats in Htt and other HEAT solenoids, and others. Much, much more work will be needed to understand how these delicate structures are coupled in turn to intrinsically disordered modules. Biophysical studies that necessarily start with truncated solenoid domains are unlikely to approximate the more dynamic lives these domains have in their full-length proteins and within the cell. Furthermore, studies focused on the biochemistry of Q/N-rich and polyQ-rich polypeptides are providing evidence of complex conformational dynamics involving multiple structural states including coiled-coils, α-helices, and β-sheets, and distinct properties promoting or disallowing multimerization and/or aggregation (30–32). Theoretical modeling has also provided some support for the ability of these conformational states to be mechanically coupled to the cell cytoskeleton (33). In this regard, a recent study has tested the complex dynamics bound to emerge in a full-length Huntingtin protein and has found evidence for the proposed intra-repeat modulatory effect on solenoid formation by its N-terminal polyQ tract (34). Additional studies in which these expanded polyQ tracts are moved either to polyQ insertion slots identified by evolutionary comparisons or to negative control slots could highlight the extent to which the concept of polyQ insertion slots is productive. If so, the identification of such slots in fast-evolving dipteran orthologs of key human proteins should be prioritized.

## Methods

Protein structures were analyzed and annotated using NCBI’s Cn3D 4.3.1 software (http://www.ncbi.nih.gov/Structure) and high resolution structures of the ankyrin repeats of human Notch1 (PDB ID: 2F8Y, MMDB: 38239) (26). The motif logo of Fig. 3b was constructed using WebLogo (http://weblogo.berkeley.edu/logo.cgi). The source sequences for dipteran Notch proteins are from the following accessions and additional analyses: AAC36153.1, AAC36151.1 (*Lucilia cuprina*), XP_011292982.1 (*Musca domestica*), XP_013109509.1 (*Stomoxys calcitrans*), XP_004535280.1 (*Ceratitis capitata*), and XP_014087559.1 (*Bactrocera oleae*). The NICD sequence of *Glossina morsitans* (Tsetse fly) is a conceptual translation of CCAG010014983. The NICD sequence of *Anopheles darlingii* is a combination of ETN64594 and a splice corrected sequence from ADMH02000947. Alignments of Notch intrinsically disordered protein regions were explored by extensive variation of parameter space using both CLUSTALW (35) and MUSCLE (36, 37) on MEGA7 (38), but these were incongruent with each other and frequently subject to polyQ alignment artifact. Computed alignments improved when the sequences were anchored and or trimmed to include only the N-terminal ARIEs slot leaders to the highly conserved C-terminal DUF3454 sequence. Alignments in their final form shown in the figures were constructed manually and represent a maximization of the number of optimal local alignments.

## ACKNOWLEDGEMENTS

I thank members of the Erives Laboratory for listening to another crazy story from their advisor at lab meeting and asking good questions. I thank members of the Kreinitz Laboratory for the same. I thank Dr. Jan Fassler for providing critical feedback on an earlier version of this manuscript.

